# Updating ACC preclinical models: characterization of two new patient-derived cell lines

**DOI:** 10.64898/2026.01.15.699554

**Authors:** Andrea Abate, Mariangela Tamburello, Claudia Bonera, Guido Alberto Massimo Tiberio, Sara Baldelli, Mattia Carini, Alessia Inglesi, Maria Scatolini, Deborah Cosentini, Marta Laganà, Marta Leporati, Pietro Luigi Poliani, Domenico Cosseddu, Duilio Brugnoni, Alfredo Berruti, Sandra Sigala

**Affiliations:** Section of Pharmacology, Department of Molecular and Translational Medicine, University of Brescia, Brescia, Italy; Surgical Clinic, Department of Clinical and Experimental Sciences, University of Brescia at ASST Spedali Civili di Brescia, Brescia, Italy; Central Clinical Laboratory, ASST Spedali Civili di Brescia, Brescia, Italy; Molecular Oncology Laboratory, Fondazione Edo ed Elvo Tempia, Ponderano, BI, Italy; Medical Oncology Unit, Department of Medical and Surgical Specialties, Radiological Sciences, and Public Health, University of Brescia at ASST Spedali Civili di Brescia, Brescia, Italy; Laboratory Medicine, AO Ordine Mauriziano, Turin, Italy; Pathology Unit, Department of Molecular and Translational Medicine, University of Brescia, Brescia at ASST Spedali Civili di Brescia, Brescia, Italy

**Keywords:** adrenocortical carcinoma, cell lines, primary cultures, experimental models, hormones secretion

## Abstract

AdrenoCortical Carcinoma (ACC) is an aggressive, rare and heterogenous malignancy, that requires diverse preclinical models. For this reason, the development of new cell lines is pivotal. Here we describe the development and characterization of two of them, SMAC-2 and SMAC-3, established from surgical specimens of ACC patients. The characterization included their mutational profiling, the evaluation of steroidogenic enzymes expression, secretory activity and the expression of steroid hormone receptors. The proliferative ability of these cells within a zebrafish embryos xenograft was also evaluated. SMAC-2 originated from a metastatic EDP-M-treated ACC in a female patient with Cushing syndrome and hyperandrogenism, while SMAC-3 derived from a male patient with a mitotane-treated local recurrence, with no sign of hypercortisolism. *TP53* was mutated in both lines. SMAC-2 cells were characterized by a pathogenic alteration on *CTTNB1* gene and a deletion of *CDKN2A* gene, while SMAC-3 on *MSH2* gene. Basal hormonal status analysis showed a cell model-specific fingerprint either in the hormonal secretion and gene and protein expression of steroid hormone receptors. SMAC-2 secreted high levels of cortisol. SMAC-3 secreted low basal level of cortisol. Mitotane displayed in both cell lines a low potency. Under the experimental conditions used, the xenografted area did not increase for both cell models. Experiments were carried out to study the stability of the two cell lines. SMAC-2 and SMAC-3 display unique molecular and functional features, expanding the repertoire of experimental ACC models and representing valuable tools for preclinical research alongside established cell lines.

## Background

Adrenocortical carcinoma (ACC) is a rare endocrine neoplasm with an annual incidence of 0.7-2.0 cases/million people per year (1, 2). Radical surgery in an experienced center is the first-line strategy for operable diseases. If there is a high risk of relapse, adjuvant therapy with the adrenolytic drug mitotane should be considered (1, 3). In about half of patients the diagnosis of ACC occurs, however, when the disease is already metastatic and not suitable for surgery. In addition, a considerable proportion of patients undergoing radical surgery recurred within two years, often with metastatic disease (1). In advanced or metastatic stages not amenable to surgery, the approach is pharmacological based on mitotane in combination with chemotherapy agents etoposide (E), doxorubicin (D), and cisplatin (C) according to EDP-M scheme (4) (2). In case of progression after EDP-M there are no standard treatment options (5, 6). The clinical presentation of ACC is heterogeneous: the disease may present with a clinically significant hormone excess or with symptoms caused by an abdominal mass (7) (8). More than half of patients with ACC have clinical hormone excess: hypercortisolism (Cushing syndrome) or mixed Cushing and virilizing syndromes. Pure androgen excess is less frequent, while estrogens or mineralocorticoid excess is very rare (9) (10) (11, 12). Excess cortisol is an independent prognostic factor and is associated with the worst prognosis (9). The pathogenesis of ACC is also heterogeneous, if most cases are sporadic, others may relate to hereditary syndromes (1) (13). Lynch syndrome, characterized by mutations on genes involved in DNA mismatch repair genes, is an example (14). In sporadic ACC, the most frequently reported somatic mutations are affecting p53 signaling, Wnt/ β-catenin pathway, and IGF2 overexpression (15) (16) (17). Metastases also have a high heterogeneity, due to additional mutations compared to primary tumors (18, 19).

To study ACC biology and identify new therapeutic options, preclinical models are essential. Taking into account the heterogeneity that characterizes the disease, different cell models are required. Since 2016 the only human adult ACC cell line available were NCI-H295 cells and its subclones (20). These cells, derived from a primary disease sensitive to mitotane, represent only a small subgroup of ACC. Patients with advanced metastatic disease and pre-treated disease are those who urgently need new effective pharmacological strategies, especially in case of progression after EDP-M. In recent years, different cell lines derived from metastatic disease have been developed (21, 22) (23). However, available cell lines are limited and do not cover the full spectrum of characteristics seen in ACC-patients.

Therefore, in this study we aim to improve the panorama of available preclinical ACC models through phenotypic and molecular characterization of two new cell models by describing their development from two patient-derived primary cultures. These cell cultures were selected for their ability to replicate, subsequently deeply characterized to propose them as new experimental cell model of human ACC, named SMAC-2 and SMAC-3 cell lines.

## Materials and methods

### Patient-derived primary culture

Primary cultures ACC172 and ACC173 (subsequently named, respectively, SMAC-2 and SMAC-3 cells lines) were established from tumor tissues of patients diagnosed with ACC and underwent surgery at ASST Spedali Civili di Brescia (Brescia, Italy). The collection of biological samples was approved by the Ethics Committee (NP1924) and informed consent was received from the patients. The primary cultures were obtained as previously described (24, 25) and both cultured in advanced D-MEM/F12 (Thermo Fisher Scientific, Waltham, MA, USA), 10% FBS (Thermo Fisher Scientific), 292 μg/mL L-glutamine (Merck Italia, Roma, Italy), 100 I.U./mL penicillin + 100 μg/mL streptomycin (Merck Italia) and 1% amphotericin B 2.5 μg/mL (Euroclone, Milano, Italy). The culture plates and multi-well plates used for SMAC-3 were treated with poly-L-lysine (final concentration: 0.01%, Merck Italia) to promote cell adhesion.

### Cell lines

Human ACC cell line NCI-H295R, human prostate cancer cell line LNCaP and human ductal carcinoma of the breast cell line T-47D were purchased from ATCC (American Type Culture Collection, Manassas, VA, USA) and cultured as indicated by the company. Cell lines were periodically tested for mycoplasma and authenticated using Short Tandem Repeats (STR) profile by BMR Genomics S.r.l. (Padova, Italy). The STR profile were analyzed also for the original tumor tissues.

### Tissue stainings

Human tissue samples of the tumors from which the cell lines were derived have been retrieved from the Institutional database of the Department of Pathology (ASST Spedali Civili di Brescia). Histological diagnosis was revised according to the most recent World Health Organization (WHO) classification of endocrine and neuroendocrine tumours (WHO classification of tumours series, 5th ed.; vol. 10). Representative sections from formalin-fixed and paraffin embedded blocks were used for routine Hematoxylin and Eosin (H&E) staining and immunohistochemistry as previously described (26). Primary antibodies are listed in **Supplementary Table 1**. Reaction was revealed by Dako EnVision+Dual Link System Peroxidase (Dako Cytomation, Carpinteria, CA, USA) followed by DAB and counterstaining with Hematoxylin. Images were acquired with an Olympus Bx60 microscope equipped with a DP70 camera (Olympus, Segrate, Italy) and CellF imaging software (Soft Imaging System GmbH; Münster, Germany).

### Hematoxylin and eosin cell culture staining

Cells were grown onto 12 mm poly-L-lysine-coated coverslips and then fixed with Immunofix® (Bio-Optica Milano, Italy) for 15 minutes at room temperature. The samples were incubated with Carazzi haematoxylin (DiaPath, Bergamo, Italy) for 5 minutes and then washed extensively to remove excess of staining. Following a 30 second incubation with eosin 1% aqueous solution (FoLAbo, Milano, Italy) was performed. Other extended washes were performed. The samples were subsequently dehydrated by exposure to an ethanol gradient (70%-100%), followed by xylene. Finally, the coverslips were mounted using DPX Mountant for histology (Merck Italia). Images were acquired using an Olympus IX51 optical microscope (Olympus) equipped with a 20x and 40x objectives.

### Cell treatment, cell viability and cell proliferation

To define the optimal treatment time for SMAC-2 and SMAC-3, the doubling time was calculated. Cells were seeded (2×10^4^ cells/well) in 4-wells and fixed with Immunofix® at different time-points. The doubling time of NCI-H295R cells, used as control for mitotane sensitivity, was previously calculated using the same protocol. Cells were incubated for 20 minutes with a 0.1% of crystal violet (Merck Italia) solution prepared in 20% methanol. After extensive washes, the staining was eluted with a 10% aqueous acid acetic solution. The absorbance at 595nm and 655nm were acquired with an EnSight Multimode Plate Reader (PerkinElmer, Waltham, MA, USA). The absorbance values were interpolated using a calibration curve to determine the cell number. The calculation of the doubling time was performed according to the ATCC method. To evaluate the effect of a treatment on cell viability and proliferation, ACC cell lines were seeded (1×10^4^ cells/well) in 96-wells plates and treated with increasing concentrations of mitotane (SMAC-2: 25 – 80µM; SMAC-3: 25 – 60µM for cell viability assay and 10 – 60µM for proliferation assay; NCI-H295R: 3 – 48µM for both cell viability and cell proliferation) and forskolin (5-25µM) for up to 96 hours. Mitotane (Selleckchem Chemicals, DBA Italia, Segrate, Milano, Italy) was dissolved in 100% dimethyl sulfoxide and stored at −20°C in 180 mM aliquots. Forskolin (FSK) (Merck Italia) was dissolved in 100 % dimethyl sulfoxide and stored at −20°C in 100 mM aliquots. Cell viability was evaluated by WST-1 assay (Merck Italia) according to while cell proliferation was evaluated by chemiluminescent BrdU-incorporation assay (Merck Italia) following the manufacturer’s instructions. Absorbance and chemiluminescence signals were detected using an EnSight Multimode Plate Reader.

### DNA and RNA extraction

DNA and total RNA were extracted from cell pellets using QIAamp® DNA Blood Mini kit (Qiagen, Hilden, Germany) and RNeasy® Mini Kit (Qiagen) respectively following the manufacturer’s instructions. During the extraction of total RNA, to eliminate any contamination from DNA, a treatment with RNase-Free DNase Set (Qiagen) was performed, according to the manufacturer protocol.

### Mutational status analysis

RNA and DNA extracted from cell cultures SMAC-2 and SMAC-3, were characterized by Next-generation Sequencing (NGS) methodology with the CE-IVD Myriapod® NGS cancer probe PLUS kit (Diatech Pharmacogenetics s.r.l., Ancona, Italy). The panel allows for the characterization of solid tumours by evaluating the main prognostic and therapeutic targets on DNA and RNA level (**Supplementary Table 2**). The panel can identify both DNA mutations (SNV, indels), copy number variants (CNV, amplification and deletions) and RNA fusion transcripts.

The DNA was quantified in Real-Time qPCR by EasyPGX® qPCR instrument 96 (Diatech Pharmacogenetics s.r.l., Ancona, Italy) and EasyPGX® Analysis Software version 4.0.15 (Diatech Pharmacogenetics s.r.l., Ancona, Italy) in order to evaluate amplifiability. A total of 60 nanograms of RNA were subjected to retrotranscription, using specific target primers, and enrichment assay. This assay allows the retro-transcription and further amplification of the cDNA. The cDNA obtained was combined with 100ng of corresponding genomic DNA. Libraries were prepared by automation using the MagnisDx NGS Prep System (Agilent Technologies, Santa Clara, California, USA). The captured libraries were then quantified by Qubit dsDNA HS Assay Kits (Invitrogen, Carlsbad, CA, USA) on Qubit 3.0 fluorometer and normalized for quantity (4nM) to ensure homogenous coverage of samples during sequencing. Finally, the libraries were pooled and sequenced in platform Mi-Seq® System (Illumina, San Diego, CA, USA). The analysis of FASTQ files were performed with the Myriapod® NGS Data Analysis Software (Diatech Pharmagenetics s.r.l, Ancona, Italy, version 5.0.11).The software aligned the sequences to the GRCh37 (hg19) reference genome (BAM file), and the analysis includes variant calling and Copy Number Variant (CNV) prediction using a bioinformatics algorithm. The reference database used for the classification of variants is ClinVar (www.ncbi.nlm.nih.gov). Variants with a minimum reliable sequencing depth of 250 clusters and Variant Allele Frequency (VAF%) ≥ 5 were evaluated. Furthermore, variants classified as benign (class 1) and likely benign (class 2) are not reported

### Gene expression

1μg of RNA extracted as described above was retrotranscribed into cDNA using the reverse transcriptase of M-MLV (Promega Italia, Milano, Italy) and Oligo(dT)15 Primer (Promega Italia) following the manufacturer’s instructions. Gene expression was evaluated by qRT-PCR (ViiA, Applied Biosystems, Milano, Italy) using SYBR Green as fluorochrome (Invitrogen, Thermo Fisher Scientific), as described in (27). The oligonucleotide sequences of the primers are listed in **Supplementary Table 3**. β-actin was used as a housekeeping gene. The difference in the threshold cycle (Ct) between the gene of interest and β-actin (ΔCt) was used as an indicator of mRNA expression.

### Immunofluorescence

Cells were grown onto 12 mm poly-L-lysine-coated coverslips and then fixed, permeabilized and blocked as previously described (26). Cells were the incubated over-night at 4°C with primary antibodies. After extensive washes, the secondary antibodies were applied at room temperature for 1 hour followed by counterstaining with Hoechst (Merck Italia) for 5 min. Primary and secondary antibodies are listed in **Supplementary Table 1**. After rinsing coverslips were mounted using FluorPreserveTM Reagent (Merck Italia) and cell staining was detected using a Zeiss LSM 900 confocal laser-scanning microscope equipped with an EC Plan-Neofluar 40x/1.30 immersion objective (Carl Zeiss S.p.A., Oberkochen, Germany). Zen blue software (Carl Zeiss S.p.A.) was used for image analysis and processing.

## Collection of conditioned supernatants

To investigate the basal secretion of steroid hormone, SMAC-2 and SMAC-3 cells were seeded in 6-well plates (1.5×106 cells/well). The following day, the culture medium was replaced with serum-free simplified medium (1mL/10^6^ cells): 1:1 (v/v) DMEM-Low Glucose (Sigma Aldrich) and Nutrient Mixture F-12 Ham (Sigma Aldrich), supplemented with 292 µg/mL of L-glutamine (Sigma Aldrich). After 24 hours, the medium was collected and frozen at -80°C. To investigate the modulation of steroid secretion the cell cultures were seeded in 6-well plates (1×10^6^ cells/well) and treated with FSK (25µM), with their respective mitotane IC_10_ values (SMAC-2 cells: 21 µM, SMAC-3 cells: 15 µM), angiotensin II (Ang II; 250nM; Sigma Aldrich) and potassium chloride (KCl; 20mM; Sigma Aldrich) for up to 4 days. The concentrations of Ang II and KCl used were selected following the ranges published in Sigala et al (28). At the end of the treatment the cells were cultured in serum-free medium for 24h as indicated above.

### Liquid Chromatography Tandem Mass Spectrometry (LC–MS/MS) Steroid Measurements

Secreted steroids concentrations were determined by two laboratories using an ultraperformance liquid chromatography tandem mass spectrometry method. Collected supernatants were purified by solid phase extraction and stable-isotope dilution was applied. For quantification, multiple-reaction monitoring mode was used. The method was a modification of the CE-IVD MassChrom® method for the quantification of steroids in serum/plasma (ChromSystems, Gräfelfing, Germany) and was validated on cell medium. Obtained concentrations were comparable between the two laboratories.

### Fish and embryos maintenance

*Danio rerio* (zebrafish) were maintained and used according to EU Directive 2010/63/EU for animal use following protocols approved by the local committee (OPBA, Organismo Per il Benessere Animale) and authorized by the Ministry of Health (authorization number 393/2017). Adult transgenic line Tg (kdrl:EGFP) and wild type zebrafish lines were maintained as described in (29). Breeding of adult male and female zebrafish was carried out through natural crosses, and embryos were collected and raised in fish water with incubation at 28.5°C until the experiments. Embryos at 24 hours post fertilization (hpf) were treated with 0.003% 1-phenyl-2-thiourea (PTU) to prevent pigmentation. After the conclusion of the experiments, the zebrafish embryos were euthanized with 400 mg/L tricaine (ethyl 3-aminobenzoate methanesulfonate salt; Merck Italia).

### ACC cell xenograft

ACC cells were exposed overnight with the vital red fluorescent dye CellTrackerTM CM-DiI Dye (Thermo-Fisher Scientific), according to the manufacturer protocol. Cells were detached with trypsin/EDTA, washed in PBS, resuspended in 50 µL of PBS, and kept at 4°C until use. Cell xenografts were performed as described in (30) with minor modifications. Briefly, zebrafish embryos at 48 hpf were dechorionated, anesthetized with 0.042 mg/mL tricaine (Sigma-Aldrich), and microinjected with the labelled ACC cells into the sub-peridermal space of the yolk sac. Microinjections were performed with a FemtoJet electronic microinjector coupled with an InjectMan N12 manipulator (Eppendorf Italia, Milano, Italy). Approximately 250 cells/4 nL were injected into each embryo; embryos were maintained in PTU/fish water in a 32°C incubator. Pictures of injected embryos were acquired using Zeiss Axiozoom V13 (Carl Zeiss AG) fluorescence microscope, equipped with Zen pro software, 2 hours (T0) and 3 days (T3) after cell injection. The tumor areas at T0 and T3 were measured using Zen 2.3 Black software (ZEISS, Jena, Germany). Representative embryos at T3 stage were fixed, embedded in low melting agarose, and acquired using a Zeiss LSM 900 confocal laser-scanning microscope (Carl Zeiss S.p.A.) equipped with a 10x objective.

### Statistical analysis

Statistical analysis was carried out using GraphPad Prism software version 10 (GraphPad Software, La Jolla, CA, USA). One-way ANOVA was used for multiple comparisons. Where appropriate, the unpaired t-test was used. Unless otherwise specified, data are expressed as mean ± Standard Deviation (SD) of at least three experiments run in triplicate. P values < 0.05 were considered statistically significant.

## Results

### Establishment of two novel ACC cell lines, SMAC-2 and SMAC-3

Two primary cultures (ACC172 and ACC173 cells) were established from two patients diagnosed with ACC and underwent surgery. ACC172 cells derived from the primary tumor mass in a metastatic ACC female patient in progression after EDP-M. ACC173 cells derived from a lymph node localized in the adrenal lodge of a local relapse of ACC in a male patient treated with mitotane. The clinical characteristics of the patients and the histological analysis of the tumor are reported in **Table 1** while representative H&E, SF-1 and Ki67 stainings on the original tumor tissues is presented in **Figure 1A**. The adrenal origin of the two established cell cultures was assessed both at mRNA level (ΔCt ± SD: ACC172 4.036 ± 0.519; ACC173 6.083 ± 1.29) and protein level (a representative immunofluorescence of SF-1 is shown in **Figure 1B)**. The two cell cultures proved to be continuously passable *in vitro* and were maintained in culture until passage 60, allowing an in-depth characterization. The active proliferation of the two cultures was confirmed by Ki67 staining (**Figure 1C**). From now onwards, ACC172 was renamed: human ACC cell line SMAC-2 and ACC173 was renamed: human ACC cell line SMAC-3. Clear field images representing the two cell lines are shown in **Figure 1D**. The cells were authenticated by STR profile analysis at passage 1, and querying the ATCC STR database (31), using both the Masters and the Tenabe algorithms, did not show any matches ≥ 56% with the STR profiles of the cell lines present in the database. To monitor the stability of the two cell models, STR analyses were periodically repeated up to 60 passages. For SMAC-2 cells, the STR profile remained constant, whereas for SMAC-3 cells, microsatellite instability was observed. STR profile analysis was also performed on the tissue samples collected during surgery.

**Figure 1.**
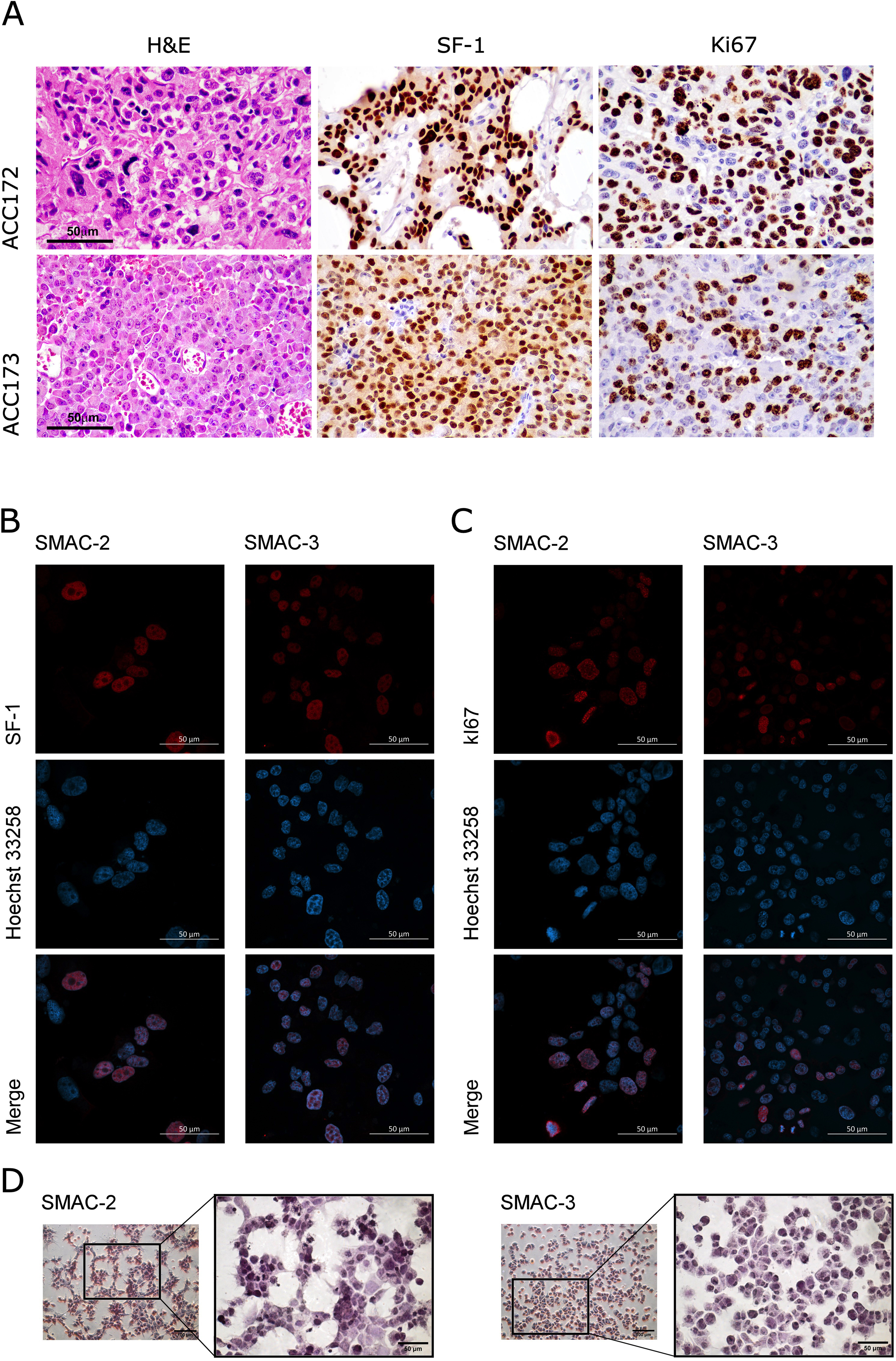
Tissues and cells characterization. (A) Histological features and expression of SF-1 and Ki67 in tumor specimens from which cell lines were derived. Upper panels show representative H&E and immunohistochemical stainings from the ACC tumor related to ACC172 cell line. Tumor show a mitotically active tumor composed by atypical cells with finely granular eosinophilic cytoplasm and pleomorphic nuclei (left panel); neoplastic cells show intense and diffuse immunoreactivity for SF1 (middle panel) and Ki67 (estimated proliferative index ranging between 15 to 30%; right panel). Lower panels show representative images from the ACC tumor related to ACC173 cell line showing a ACC tumor with solid growth composed by epithelioid cells with eosinophilic cytoplasm and frequent nuclear atypia with prominent nucleoli (left panel); tumor cells were immunoreactive for SF1 (middle panel) and Ki67 (estimated proliferative index ranging between 12 to 32%; right panel). All images are from ×40 original magnification. SF-1 (B) and Ki67 (C) immunofluorescences in ACC172 (SMAC-2) and ACC173 (SMAC-3) cells. Pictures were acquired with a Zeiss LSM 900 confocal laser-scanning microscope equipped with a 40x objective. The scale bar of 50µm is automatically inserted by the software ZEN Blue. (D) H&E staining for ACC172 (SMAC-2) and ACC173 (SMAC-3). Images were acquired using an Olympus IX51 optical microscope equipped with 20x (small panel) and 40x (big panel) objectives. The scale bars of 100µm and 50µm are inserted with ImageJ software (Nation Institute of Health. Bethesda, MD, USA).

**Table 1.**
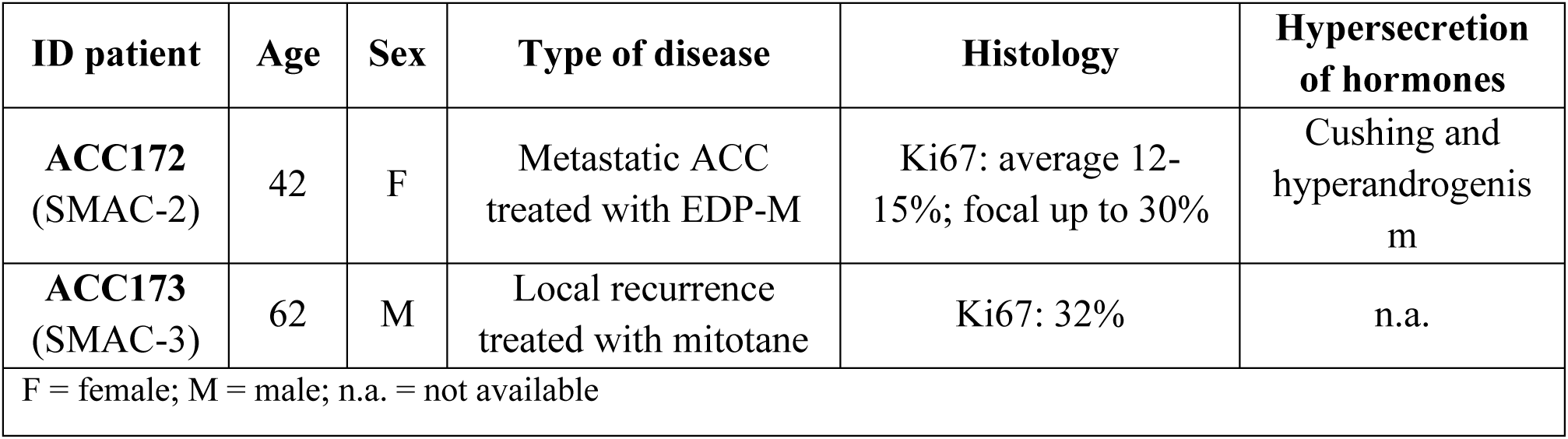
Characteristics of patient and disease.

The results of the STR profiles analyses are reported in **Table 2**.

**Table 2.**
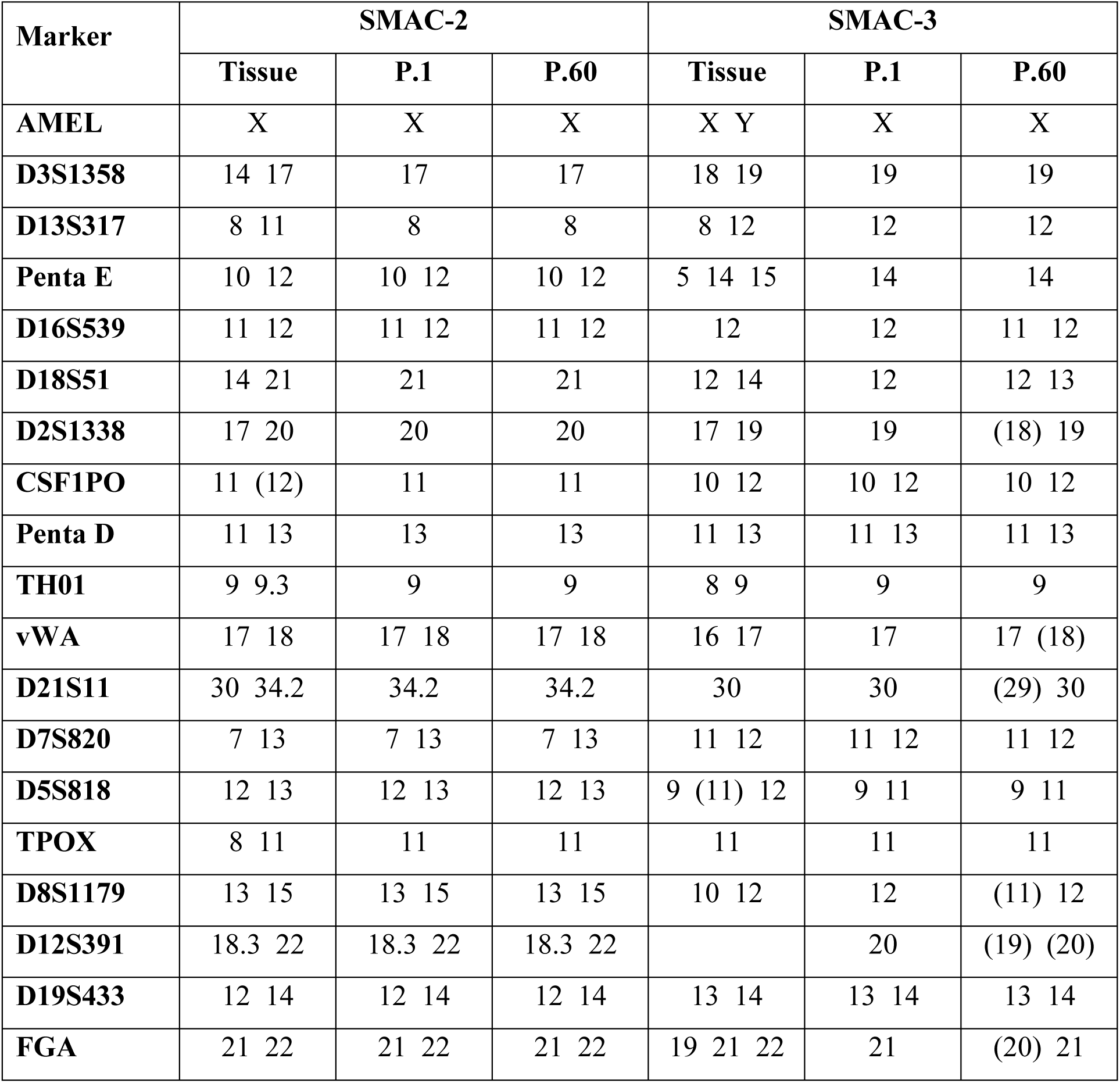
STR profiles.

### Characterization of mutational status

Taking advantage of a panel used for the characterization of solid tumors, which includes some of the known ACC driver gene (17) (15), the mutational status of the two cell lines was evaluated. Among the mutations identified for SMAC-2 cells, a WNT/β-catenin pathway mutation was detected as a frameshift mutation due to deletion of bases 116_138 in the *CTNNB1* gene. Furthermore, a complete deletion of the *CDKN2A* gene, encoding the p16 protein belonging to the cyclin-dependent kinase inhibitor family (CDKI), and a partial deletion of the *TP53* gene were also detected. In SMAC-3, a pathogenic mutation of the *TP53* gene and a mutation with a possible role of pathogenicity in the *NF1* gene were reported. **Table 3** shown the full list of mutations detected by the panel for both cell models. Considering the high interest of the WNT/β-catenin pathway in the in ACC, also as a possible pharmacological target (32), intracellular localization of β-catenin has been evaluated (**Figure 2**). In SMAC-2 cells, β-catenin appears to be active in basal conditions with a signal spread throughout the cell and markedly in the nucleus analogous to that of NCI-H295R cells (**Supplementary Figure 1**). For SMAC-3 cells, protein localization is predominantly confined to the cytoplasmic membrane. This result suggests that, under basal conditions, there is no nuclear over-activity directly induced by β-catenin.The SMAC-3 cell line is derived from a patient with a known germline mutation on the *MSH2* gene, with a predisposition to the phenotype of the Lynch syndrome. SMAC-3 cells confirmed the mutational status of the *MSH2* gene, as a Loss of Heterozygosity (LOH) at exon 7, through Sanger sequencing (**Supplementary Materials and Methods**).

**Figure 2.**
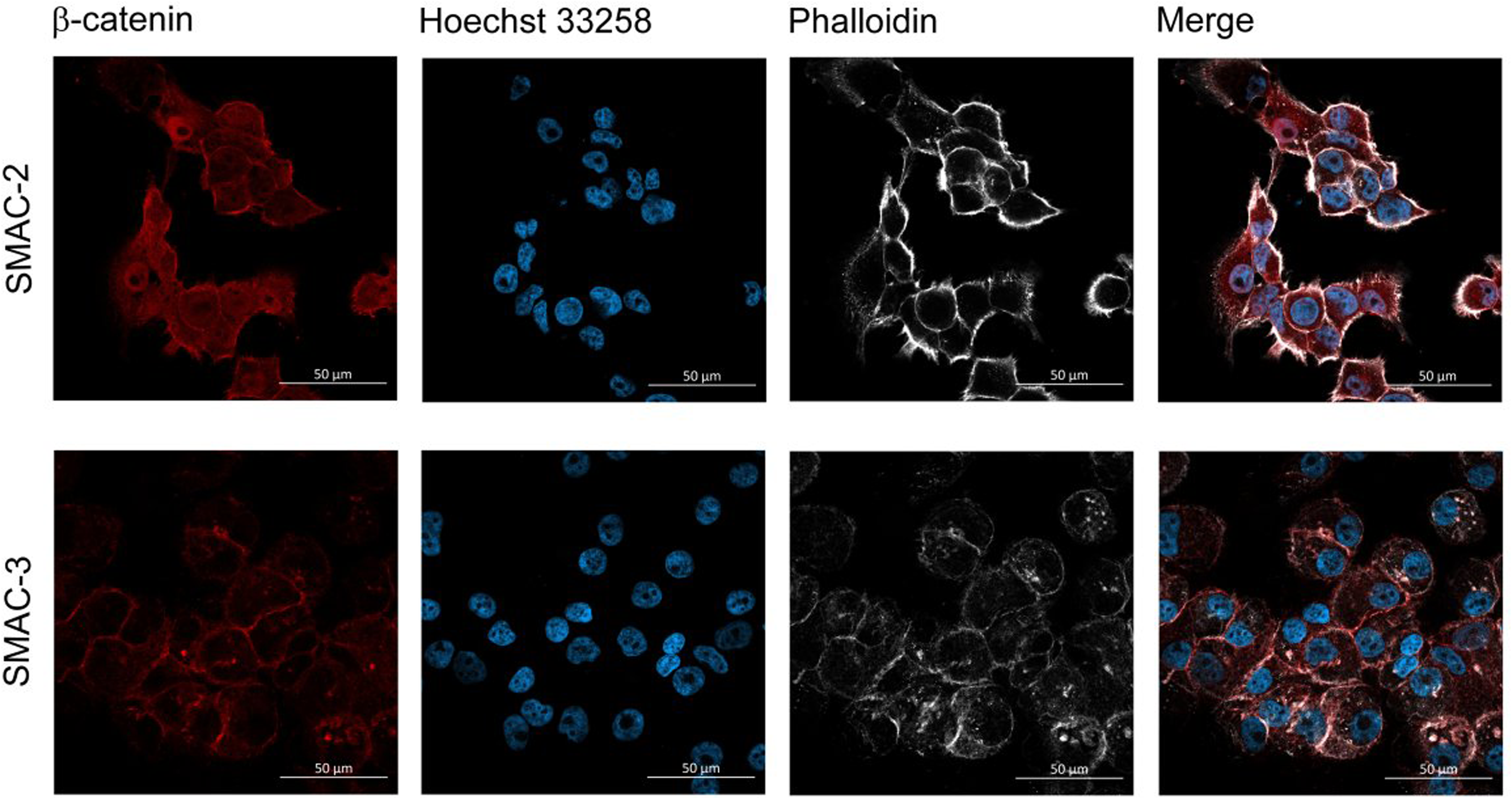
β-catenin intracellular localisation. β-catenin immunofluorescences in SMAC-2 and SMAC-3 cells. Pictures were acquired with a 40x objective. The scale bar of 50µm is automatically inserted by the software ZEN Blue.

**Table 3.**
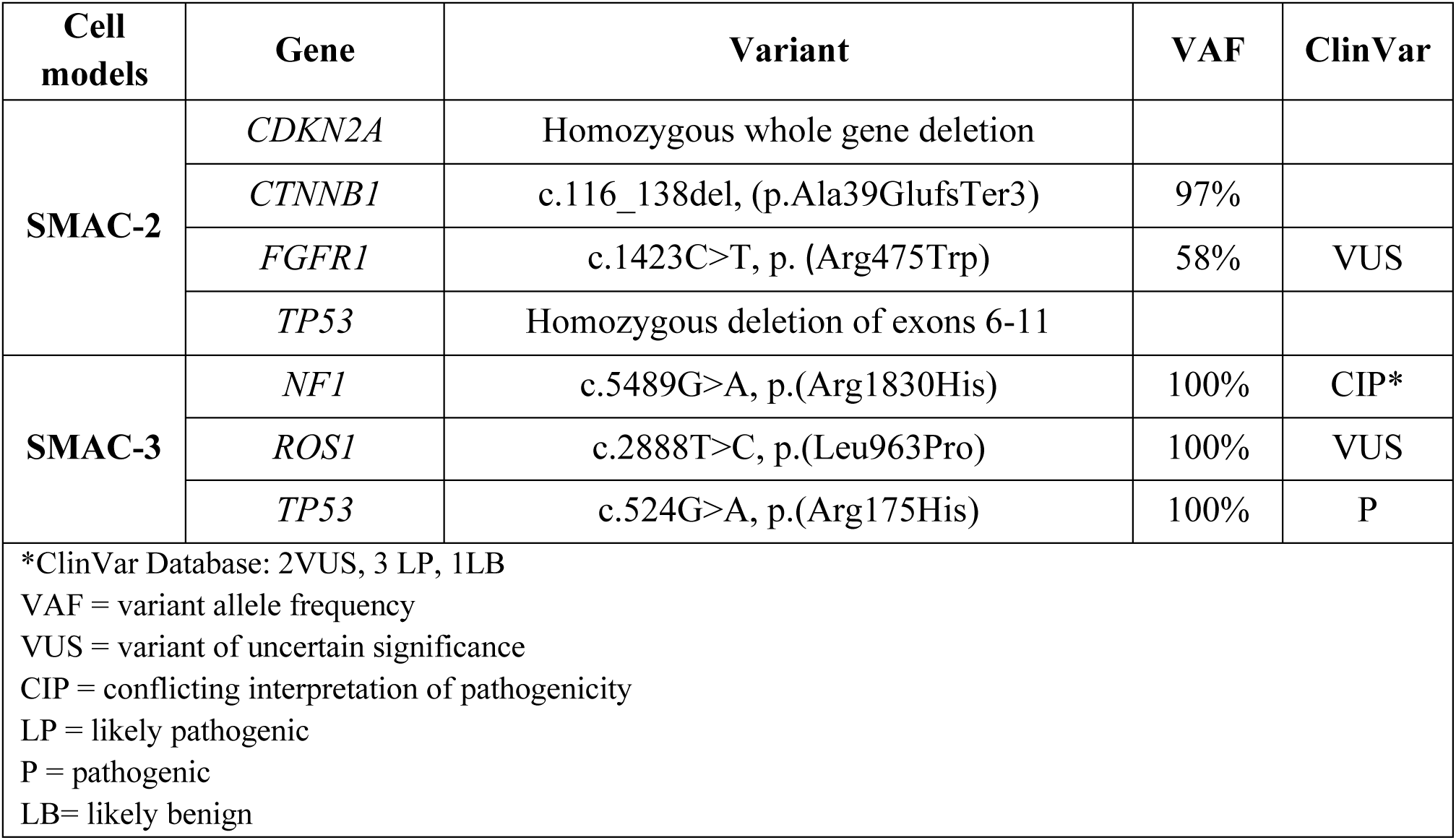
Detected mutations by the panel.

### Sensitivity to mitotane

Mitotane is the reference drug in ACC pharmacotherapy, alone or in combination with chemotherapy (2). The *in vitro* sensitivity to mitotane of the cell models was evaluated as a reduction of cell viability and proliferation after 96 hours of exposure. The treatment duration was chosen based on the calculated doubling-time (DT: SMAC-2 = 79 hours; SMAC-3 = 81 hours), to cover at least one complete duplication of the cell population. Exposure to increasing concentration of mitotane led to a concentration-dependent reduction of SMAC-2 and SMAC-3 cell viability (Emax ± SD: SMAC-2 = 48.0% ± 4.5%; SMAC-3 = 66.7% ± 9.0%) and cell proliferation (Emax ± SD: SMAC-2 = 95.3% ± 1.8%; SMAC-3 = 92.9% ± 2.7%). Sigmoidal concentration-response function was applied to generate the concentration-response curves shown in **Figure 3**. The effect of treatment on both viability and proliferation has been reassessed at high passages (P>30), showing the maintenance of the low sensitivity to mitotane (**Supplementary Figure 2**). As a positive control, the NCI-H295R cell line (DT: 52 hours), a well-known mitotane-sensitive model (33), was used. Exposure to increasing mitotane concentrations for 96 hours led to a concentration-dependent reduction in both cell viability (Emax ± SD: 100% ± 0%) and cell proliferation (Emax ± SD: 100% ± 0%). The optimal concentration ranges, reported in the Materials and Methods section, were identified based on preliminary experiments (**Supplementary Figure 3**).

**Figure 3.**
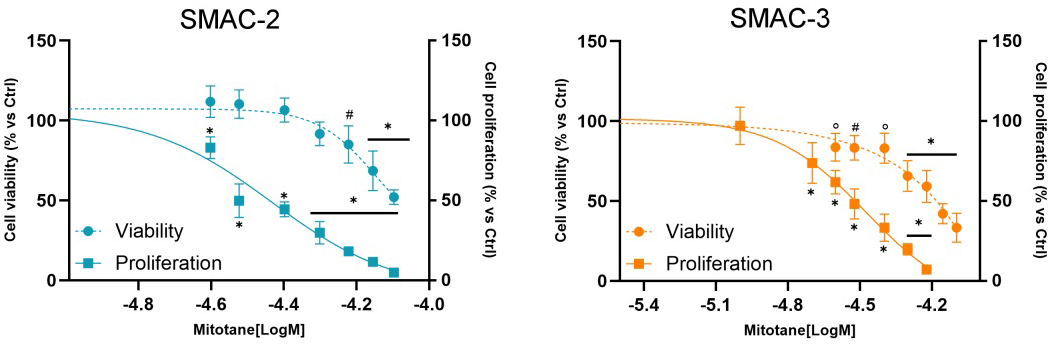
Effect of mitotane on cell viability and cell proliferation. Cells were treated with increasing concentrations of mitotane for 96 hours. Cell viability (dashed line) was analysed by WST-1 assay while cell proliferation (solid line) was evaluated with a BrdU-incorporation assay. Experiments were performed at passages ≤ 30. Results are expressed as percent of viable or proliferative cells vs untreated cell ± SD. # p <0.01; ° p < 0.001; * p < 0.0001 vs control.

### Characterization of hormonal secretion

To evaluate the steroidogenic activity, the expression of mRNA encoding for enzymes involved in steroidogenesis was firstly evaluated. qRT-PCR results suggested steroidogenic activity for both cell lines (**Figure 4**). The mRNA expression of steroidogenesis enzymes was also assessed in NCI-H295R as control (**Supplementary Figure 4**). An abundant expression of the CYP11A1 enzyme (2^-ΔCt^ ± SD SMAC-2: 6.49×10^-2^ ± 2.40×10^-2^; SMAC-3: 1.17×10^-1^ ± 3.20 x10^-2^), catalyzing the limiting step in the biosynthesis of steroid hormones, as well as the CYP17A1 enzyme (2^-ΔCt^ ± SD SMAC-2: 9.52×10^-2^ ± 4.60×10^-2^; SMAC-3: 5.70×10^-1^ ± 1.39 x10^-1^) were observed at gene level. CYP17A1 has both 17alpha-hydroxylase activity, which is necessary for the synthesis of cortisol, and 17,20-lyase activity which is essential for the synthesis of precursors of sexual hormones (34). The expression of CYP19A1 (aromatase) was poorly detectable.

**Figure 4.**
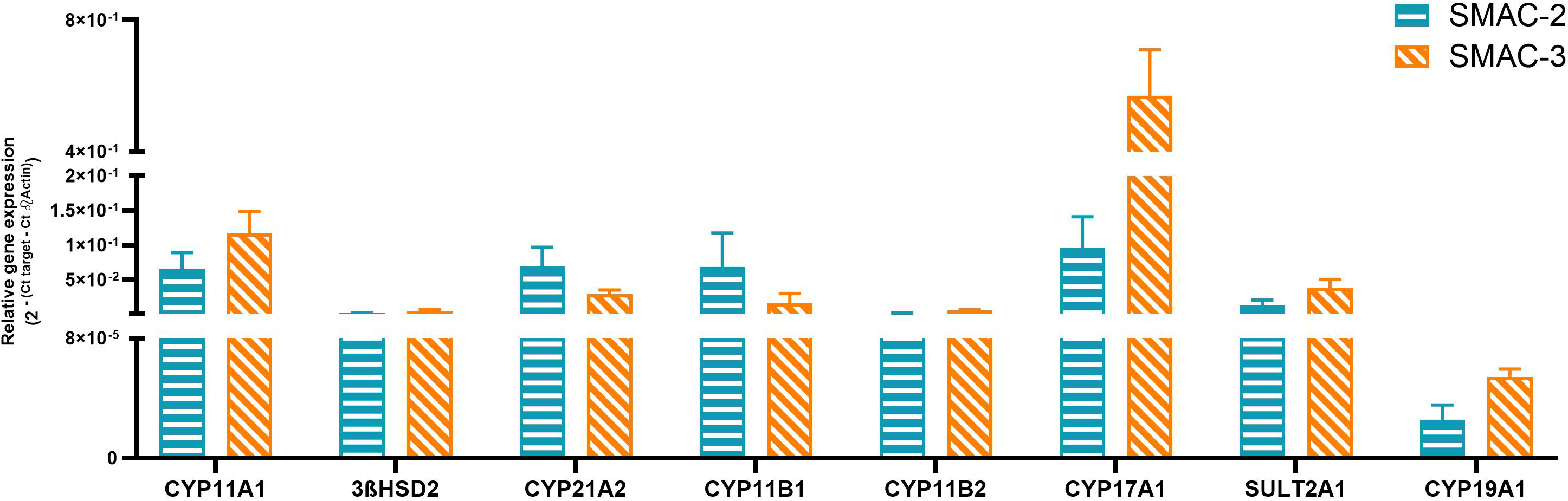
Gene expression of steroidogenic enzymes in SMAC-2 and SMAC-3 cells. Gene expressions were measured by q-RT-PCR using SYBR Green as fluorochrome. Results are shown as relative gene expression with β-actin as housekeeping gene ± SD.

To obtain a functional data on the steroidogenic activity, the basal secretion of steroids has been evaluated (**Table 4**).

**Table 4.**
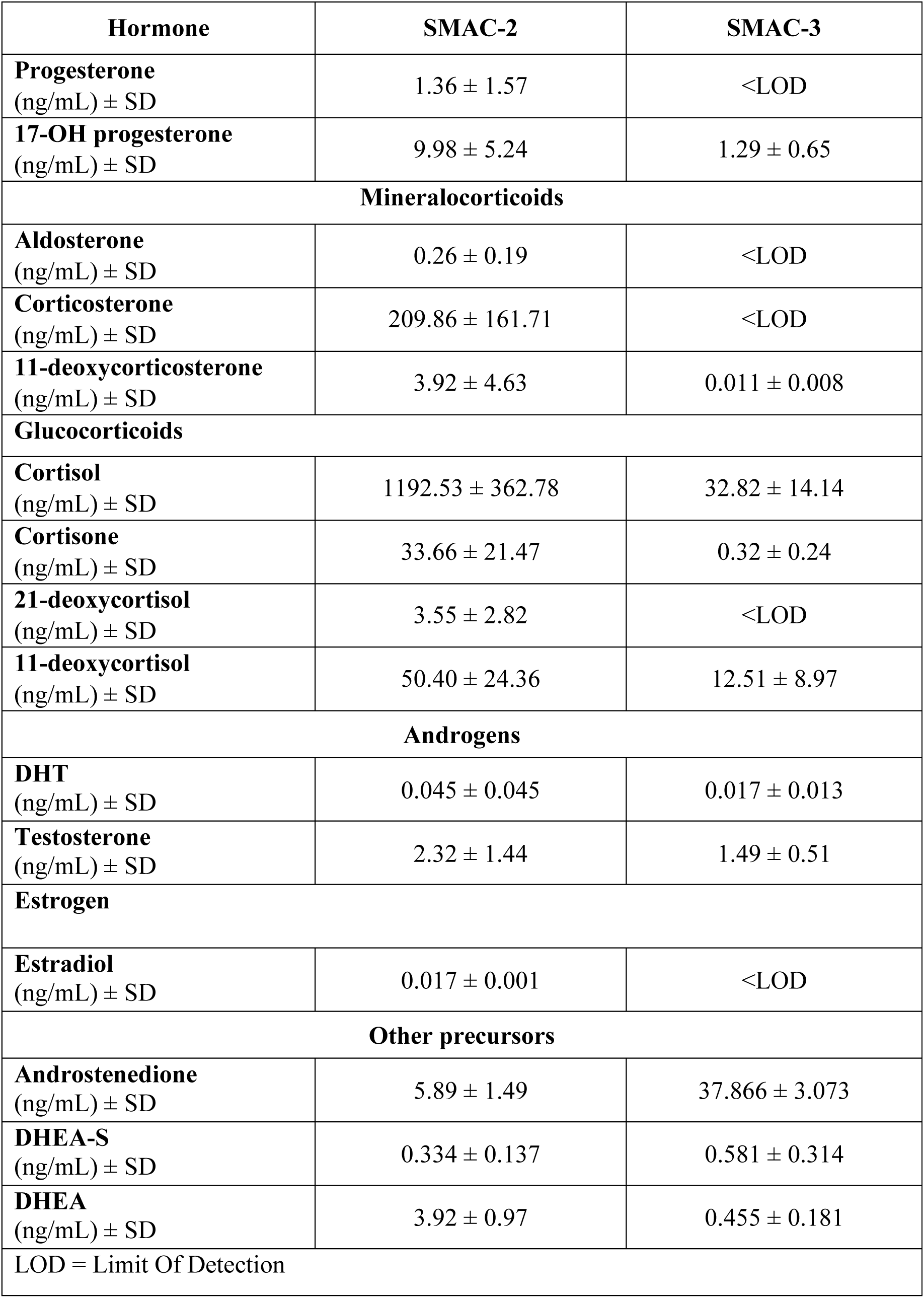
Steroid hormones profile.

In line with the gene expression data, both cell lines demonstrated an active steroidogenesis, although with model-specific features. SMAC-2 cells, according with the patient’s clinical background (Cushing’s syndrome), secreted high amount of cortisol. Abundant production of androstenedione, a common precursor of androgens and estrogens, was observed in SMAC-3. Moreover, although the original patient did not exhibit clinically significant cortisol hypersecretion, SMAC-3 cells were found to secrete cortisol. Considering the role of cortisol in the progression and prognosis of ACC (9), the effect of mitotane and forskolin on cortisol secretion was evaluated. Ninety-six hours treatment with low concentrations of mitotane (IC_10_ of cell viability: SMAC-2 cells 21 µM, SMAC-3 cells: 15 µM) led to a statically significant reduction of cortisol secretion in both models (Δ ng cortisol/mL/10^6^ cells ± SEM. SMAC-2: Δ = -1462 ± 417.7, p = 0.0387; SMAC-3: Δ = -56.96 ± 10.30, p = 0.0312) (**Figure 5A**). In contrast, exposure to 25µM of forskolin for 96 hors did not induce a statically significant increase in cortisol secretion (Δ ng cortisol/mL/10^6^ cells ± SEM. SMAC-2: Δ = 361.9 ± 342.0, p = 0.3308; SMAC-3: Δ = 7.62 ± 16.36, p = 0.6657) (**Figure 5B**). The effect of forskolin treatment on cell viability is reported in **Supplementary Figure 5**. Interestingly, as the passage progressed, the amount of cortisol secreted by SMAC-3 increased. The difference between the concentration of cortisol in the culture medium until passage 30 and after passage 30 is significantly different (**Supplementary Figure 6A**). SMAC-2, on the other hand, showed stable secretive activity. The effect of treatment with mitotane and forskolin on cortisol secretion was, therefore, also evaluated at passages greater than to 30. Results show (**Supplementary Figure 6B**) that even at higher passages, mitotane maintains an inhibitory effect on cortisol secretion, whereas the effect of forskolin remains non-significant, although a trend toward increased secretion is observed. The effect of other stimuli was evaluated on the steroidogenic activity of SMAC-2 and SMAC-3 cells. Specifically, the effect of angiotensin II and potassium on aldosterone production was assessed. As reported in Table 4, SMAC-2 cells are capable of secreting aldosterone, albeit at low levels, whereas SMAC-3 cells showed no detectable aldosterone secretion. Both stimuli did not modify the basal hormone secretion (**Supplementary Figure 7**).

**Figure 5.**
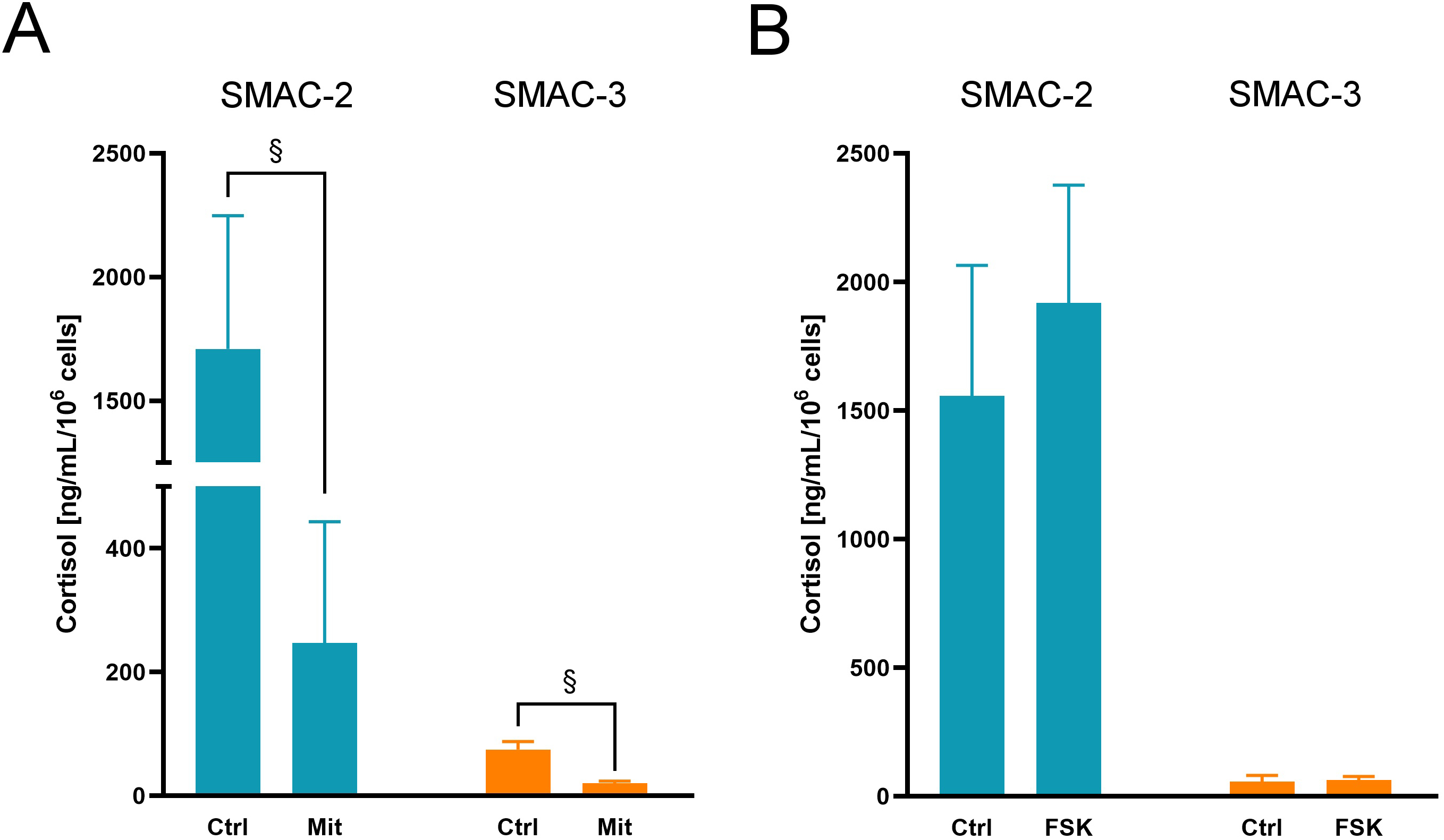
Effect of mitotane (A) and forskolin (B) on cortisol secretion. Experiments were performed at passages ≤ 30. Results are shown as ng/mL/10^6^ cells of cortisol vs untreated cell ± SD. $ p < 0.05 vs control.

The characterization was completed by evaluating the expression of steroid hormone intracellular receptors both at mRNA (**Figure 6A, C, E**) and protein level (**Figure 6B, D, F) (Supplementary Figure 8 for positive controls**). Results demonstrated the expression of the glucocorticoid receptor (GCR) in both models, while a poor expression of androgen receptor (AR) was observed. SMAC-2 also expressed detectable levels of the nuclear progesterone receptor (PgR). Additionally, the Pg membrane receptors were also investigated at mRNA level (**Supplementary Figure 9**), revealing a heterogeneous expression. Finally, the mRNA coding for the ACTH receptor MC2R was measured, showing for both models a lack/very poor expression (ΔCt ± SD: SMAC-2 14.204 ± 1.69; SMAC-3 14.671 ± 0.754).

**Figure 6.**
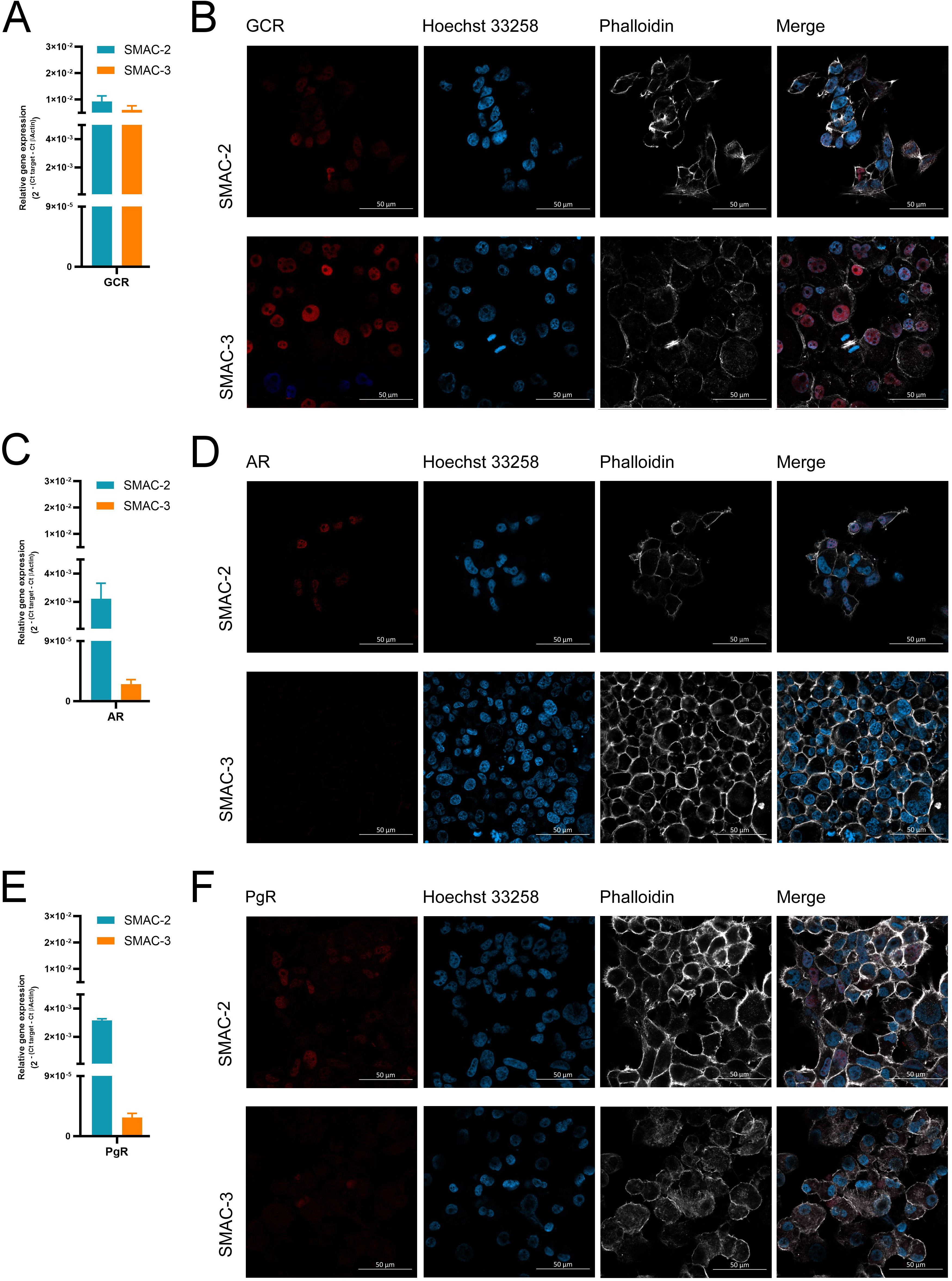
Gene (A, C, E) and protein expression (B, D, F) of intracellular steroid hormones receptors. Glucocorticoids receptor (GCR), androgens receptor (AR) and progesterone receptor (PgR) were evaluated. Gene expressions were measured by q-RT-PCR using SYBR Green as fluorochrome. Results are shown as relative gene expression with β-actin as housekeeping gene ± SD. Protein expression was investigated by immunofluorescence. Pictures were acquired with a Zeiss LSM 900 confocal laser-scanning microscope equipped with a 40x objective. The scale bar of 50µm is automatically inserted by the software ZEN Blue.

### Zebrafish-embryos xenograft

We recently demonstrated that zebrafish embryos xenografted with human ACC cells can be a fast and low-cost pharmacological tool for drug-screening (35) (36) (30). SMAC-2 and SMAC-3 were injected into zebrafish larvae and after 3 days (T3) of incubation the xenograft areas were compared with the post-injection (T0) measured areas. For both cell models there were no significant growth of the areas (Area ± SD. SMAC-2: T0 = 35460µm^2^ ± 8345µm^2^; T3 36309µm^2^ ± 14694µm^2^. SMAC-3: T0 = 26078µm^2^ ± 9037µm^2^; T3 28882µm^2^ ± 11148µm^2^). Representative images of zebrafish embryos injected with SMAC-3 are reported in **Figure 7** and **Supplementary Figure 10**. Finally, no metastasis formation was observed for both models.

**Figure 7.**
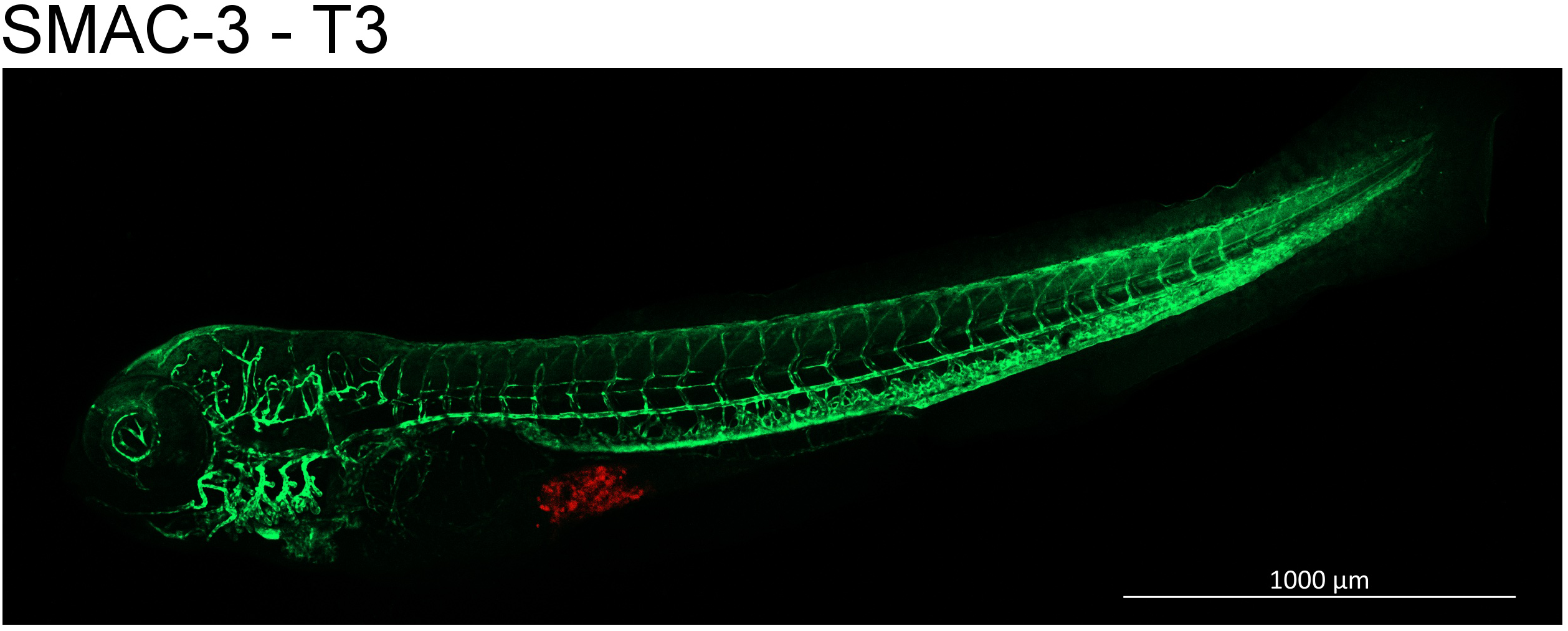
SMAC-3 cells xenograft in zebrafish embryos. Representative image of a 3 days (T3) post cells injection. SMAC-3 cells were labeled with a red fluorescent lipophilic dye while the embryo endothelium was labeled with a green fluorescent protein reporter driven by the kdrl promoter. Images were acquired using a Zeiss LSM 900 confocal laser-scanning microscope equipped with a 10x objective. The scale bar of 1000µm is automatically inserted by the software ZEN Blue.

## Discussion

There is no single, universally accepted definition of translational research. Indeed, it is a multi-step process aimed at bridging the gap between basic scientific research and clinical application. The goal of the translational research is to facilitate the process of transitioning laboratory-derived information into clinical practice (37). Among the tools of translational research, experimental cell models are certainly fundamental for studying tumor biology and for the identification of new therapeutic strategies (38) (39).

ACC is a rare and aggressive endocrine malignancy, characterized by significant morphological, molecular, and clinical heterogeneity (18) (40) (15, 19). Preclinical-translational research, therefore, requires a substantial number of cell models to recapitulate this heterogeneity. Until 2016, the only globally recognized ACC cell line was the NCI-H295R cell line and its subclones, derived from an untreated primary ACC carcinoma (20). Although this cell line has significantly contributed to the advancement of knowledge on ACC, it has the limitation of representing only a subset of patients with this malignancy. In the past decade, several ACC cell lines, both primary and metastatic in origin, have been developed (21, 22) (23). The data in this study describe the development of two new ACC cell lines, named SMAC-2 and SMAC-3. SMAC-2 derived from the primary tumor mass in a metastatic ACC female patient in progression after EDP-M while SMAC-3 was established from a lymph node localized in the adrenal lodge of a local relapse of ACC in a male patient treated with mitotane. Although not yet common in most preclinical studies based on cell cultures, it is important to highlight the gender of the patient. Indeed, an increasing amount of evidence indicates that the “sex” of the cell lines can influence the response to various stimuli (41) (42).

An in-depth characterization of the genetic landscape of the two primary cultures allowed the detection of models-specific characteristic mutations. Some of the alterations affect genes frequently mutated in ACC, such as *CTNNB1* and *TP53*. The β-catenin coding gene *CTNNB1* mutation detected in SMAC-2 has previously been described as a somatic mutation in an aldosterone-producing adrenal adenoma (Aldosterone-Producing Adenoma, APA) (43). Moreover, the same mutation is reported in the public TCGA (The Cancer Genome Atlas Program) database in a sample from a female ACC patient (44). This mutation is located near other known mutations of a gain-of-function nature (43). The similarity with the β-catenin phenotype expressed by NCI-H295R, which harbors an activating mutation on the CTNNB1 gene (45), allows us to hypothesize an activating effect for this mutation as well.

Interestingly, in SMAC-2, a homozygous deletion of 6 exons (exon 6 to 11) in *TP53* determinates the loss of TP53 DNA-binding domain. In addition, the functionality of the p53 pathway is further likely compromised due to the homozygous deletion of the whole tumor suppressor gene *CDKN2A*, which encodes the p16^INK4A^ and p14^ARF^ proteins. p14^ARF^ is part of the p14^ARF^-MDM2-p53 axis, determining p53 stabilization (46) (47) (48) (49). Inactivating mutations of *CDKN2A* are described as a negative prognostic factor in various malignancies (50, 51) and have been identified in several ACC samples (15). Recently, a case of a patient with adrenal hyperplasia developing ACC has been reported. Interestingly, mutational analysis revealed a heterozygous deletion of *CDKN2A* in the non-tumoral hyperplastic tissue and a homozygous deletion in the ACC, reinforcing the role of this mutation in neoplastic transformation (52). The *TP53* mutation found in SMAC-3, on the other hand, has a well-established pathogenic significance (53). Finally, the microsatellite instability observed in SMAC-3 is consistent with the mutation in *MSH2*, which encodes a component of the DNA mismatch repair system (54) (55).

Results about sensitivity to mitotane treatment are consistent with the origin of the cells, derived from patients with disease progression or relapse during or after mitotane therapy. Mitotane maintained a concentration-dependent effect, although effective concentrations were found to be high. Cell models that are partially/weakly sensitive to mitotane could be useful as preclinical tools for evaluating combination treatment strategies aimed at enhancing/restoring sensitivity to mitotane.

The evaluation of the steroid hormone secretion profile, together with the results related to the gene expression of enzymes involved in steroidogenesis, allows to outline a general overview of the steroidogenic activity in both cell models. Specifically, the results indicate that steroidogenesis is active in both models, leading to distinct basal secretion of steroid hormones. Regarding the differential secretion of glucocorticoids, and in particular cortisol, the findings are consistent with the clinical presentation of the two originating patients: Cushing’s syndrome for SMAC-2 and no evidence of hypercortisolism for SMAC-3. However, it should be noted that the absence of clinically relevant cortisol hypersecretion in the patient does not imply that the tumor cells are not able to secrete this hormone (56). It is interesting to note that, despite these cells being weakly sensitive to mitotane in terms of antiproliferative-cytotoxic effect, treatment at low concentrations led to a significant reduction in cortisol secretion. Taken together, these findings not only highlights e the complex mechanism of action of mitotane, (57) but also indicate that even in contexts where this compound does not exert a marked effect on the cell’s ability to survive/proliferate, it may still prove beneficial in the pharmacological management of the disease. In SMAC-3, it was observed that as the passages progressed, cortisol secretion increased. In the context of a cell line with a mutation in the DNA mismatch repair system and thus more prone to mutations, this result is not surprising. Furthermore, the *in vitro* secretory activity of ACC cells is known to depend on several factors, including the passage number (58). The variation in cortisol secretion, along with microsatellite instability, highlights the need for genetically unstable cell models such as SMAC-3, but also for more stable models to pay attention to cell culture passage number. These considerations emphasize the importance, in translational preclinical research, of performing experiments within a defined number of cell passages. Clearly reporting this information in the Materials and Methods section, alongside other critical experimental parameters, is essential to enhance the reproducibility and reliability of results.

The assessment of steroid hormone receptor expression reveals heterogeneous expression between the two models. We have previously demonstrated a potential role of progesterone as a cytotoxic agent in preclinical models of ACC (59) (36, 60). Moreover, testosterone has shown strong tumor-inhibitory properties both *in vitro* and *in vivo* (61). Specifically, knowledge of the expression levels of progesterone and androgen receptors is useful when considering the use of a cell line for evaluating the effects of endocrine treatment.

Finally, the data obtained from xenografted zebrafish embryos, although seemingly inconsistent with the previously demonstrated proliferative activity of these cell lines, should be interpreted in light of the specific experimental conditions of the xenograft model, particularly the lower incubation temperature (32°C) and the inherently slow doubling times of the two cell lines, which exceed 72 hours, the maximum duration for which zebrafish embryos can be maintained. These findings suggest that the zebrafish embryo xenograft may not be the most suitable in vivo model for evaluating the effects of pharmacological treatments on these specific cell lines.

Taken together, results presented here allowed us to present two new experimental cell models to support preclinical research in the ACC field. Indeed, given the high heterogeneity of the tumor, in addition to the currently available cell lines, SMAC-2 and SMAC-3 cell lines, with their peculiar biological and genetic traits, could help to contribute to shed a light in the complexity of this disease.

## Declarations

### Ethics approval and consent to participate

the collection of biological samples was approved by the Ethics Committee “Comitato Etico della Provincia di Brescia” (code of the study: NP1924) and informed consent was received from the patients. The facility of *Danio rerio* at University of Brescia was authorized by the Ministry of Health (authorization number 393/2017).

### Declarations of interest

the Authors declare that they have no conflict of interest.

### Fundings

This work was supported by a projected granted by AIRC (Associazione Italiana per la Ricerca sul Cancro) – IG 2022 (ID 27233, PI SS); IG 2019 (ID 23009, PI AB).

### Authors’ contributions

SS and AA conceptualized the study. GAMT operated the patients. DeC and MLa provided clinical information of the patients. AA, CB and MT performed the cell culture establishment, maintenance and treatments; MS and AI carried out the molecular genetic studies; CB and MT participated in the gene expression analysis; AA and CB performed the immunofluorescence; SB, MC and MLe carried out the steroid analysis; MT and AA carried out the experiment with zebrafish embryos. AA and CB performed the statistical analysis and wrote the manuscript. SS, AB, DoC, DB, MS, GAMT supervised the project. SS and AB provided the founding.

## Acknowledgements

We thank the LC-MS/MS technicians (Antonella, Danila, Federica, Luisella and Michael) for their excellent technical support.

## Notes

### Competing Interest Statement

The authors have declared no competing interest.

