## Supplementary Figure or Supplementary table for "Updating ACC preclinical models: characterization of two new patient-derived cell lines"

### SUPPLEMENTARY MATERIALS AND METHODS

#### Sanger Sequencing

Sanger sequencing was performed following the laboratory's standard protocol. Briefly, genomic DNA (50ng) was amplified through a polymerase chain reaction (PCR) using the AmpliTaq Gold 360 DNA Polymerase (Applied Biosystems, Foster City, CA, USA) using conditions reported in **Supplementary Table 4**. Primers targeting *MSH2* were designed with Primer3plus® (version: 3.3.0), and the primer sequences are shown in **Supplementary Table 5**. PCR products were visualized on a 2% agarose gel prepared with 1× Tris-acetate-EDTA (TAE) buffer and stained with 4.5µL of SYBR™ Safe DNA Gel Stain (Invitrogen™, Waltham, Massachusetts, USA). Subsequently, amplicons were purified using Exonuclease I and Shrimp Alkaline Phosphatase (ExoSAP, Applied Biosystems™, Foster City, CA, USA), and sequenced with the Big Dye Terminator V3.1 Cycle Sequencing Kit (Applied Biosystems™, Foster City, CA, USA). Then, sequenced products were purified using the Big Dye X Terminator Purification kit (Applied Biosystems™) according to the manufacturer's instructions. Sequencing was performed on the 3500 Dx Genetic Sequencer (Thermo Fisher Scientific, Waltham, Massachusetts, USA), and electropherograms were analyzed using Chromas v.2.6.6 and BLAST on NCBI.

#### SUPPLEMENTARY FIGURES

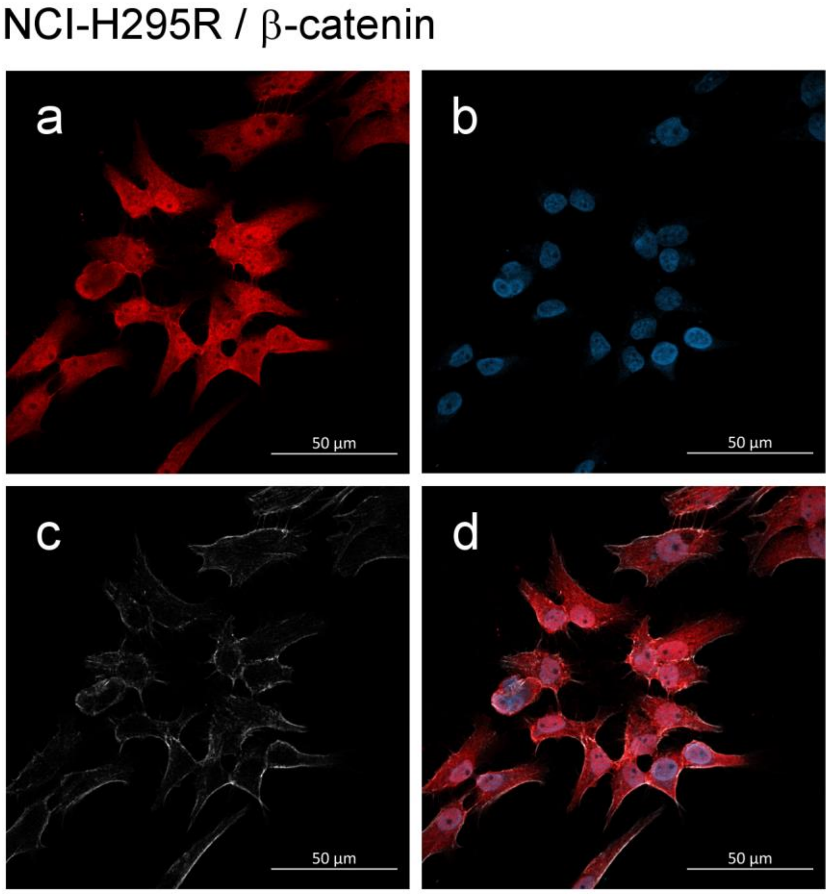


**Supplementary Figure 1. β-catenin intracellular localization in NCI-H295R cells.** (a) β-catenin, (b) Hoechst 33258, (c) phalloidin and (d) merge. Images were acquired with a 40x objective. The scale bar of 50µm is automatically inserted by the software ZEN Blue.

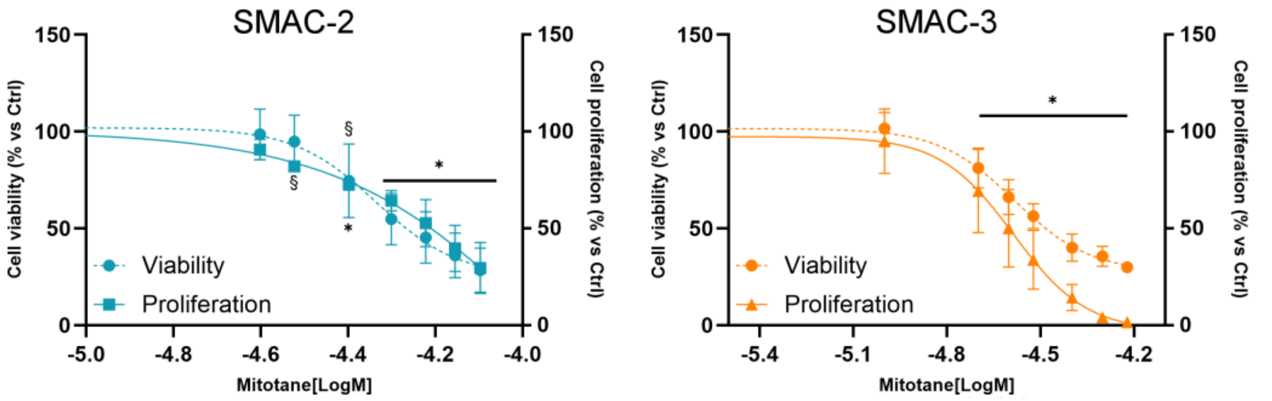


**Supplementary Figure 2. Effect of mitotane on cell viability and cell proliferation at high passages.** Cells were treated with increasing concentrations of mitotane for 96 hours. Cell viability (dashed line) was analysed by WST-1 assay while cell proliferation (solid line) was evaluated with a BrdU-incorporation assay.

Experiments were performed at passages > 30. Results are expressed as percent of viable or proliferative cells vs untreated cell ± SD. $ p < 0.05; * p < 0.0001 vs control.

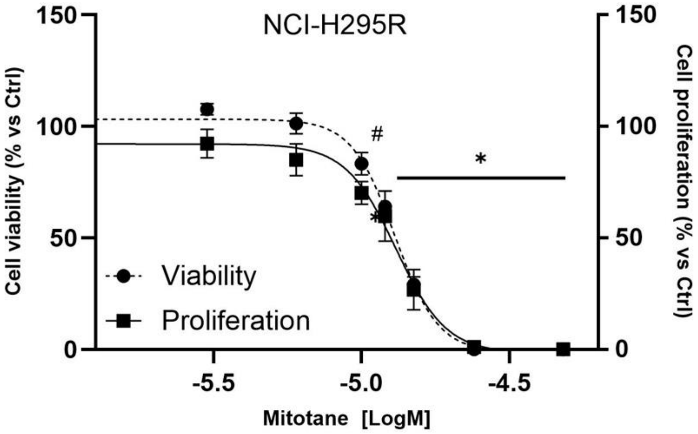


**Supplementary Figure 3. Effect of mitotane on cell viability and cell proliferation on NCI-H295R cells.** Cells were treated with increasing concentrations of mitotane for 96 hours. Cell viability (dashed line) was analysed by WST-1 assay while cell proliferation (solid line) was evaluated with a BrdU-incorporation assay. Results are expressed as percent of viable or proliferative cells vs untreated cell ± SD. # p < 0.01; * p < 0.0001 vs control.

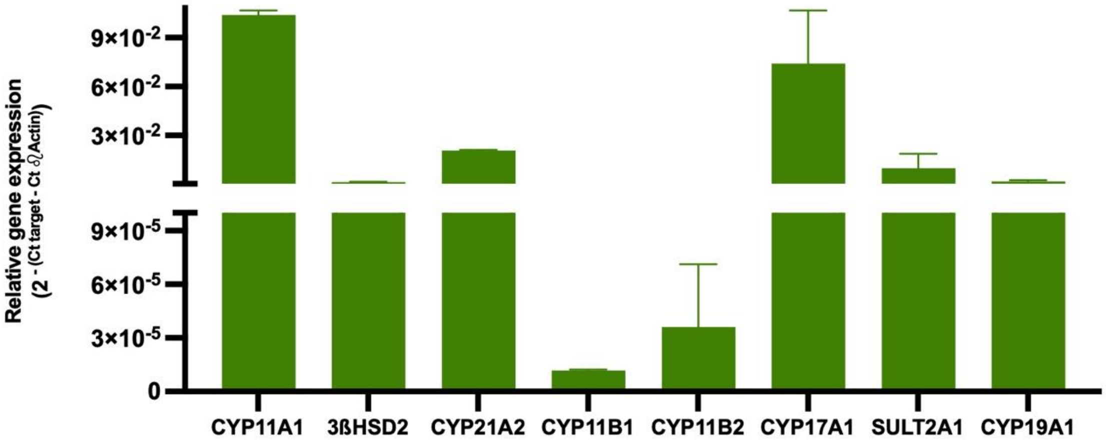


**Supplementary Figure 4. Gene expression of steroidogenic enzymes in NCI-H295R cells.** Gene expressions were measured by q-RT-PCR using SYBR Green as fluorochrome. Results are shown as relative gene expression with β-actin as housekeeping gene ± SD.

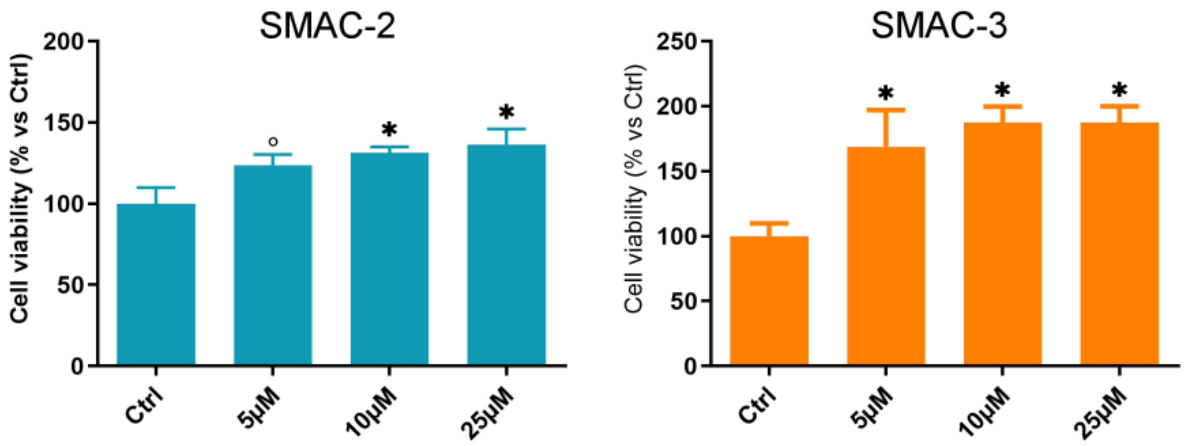


**Supplementary Figure 5. Effect forskolin on cell viability.** Cells were treated with increasing concentrations of forskolin for 96 hours. Cell viability was analyzed by WST-1 assay. Results are expressed as percent of viable vs untreated cell ± SD. ° $p < 0.001$; * $p < 0.0001$ vs control.

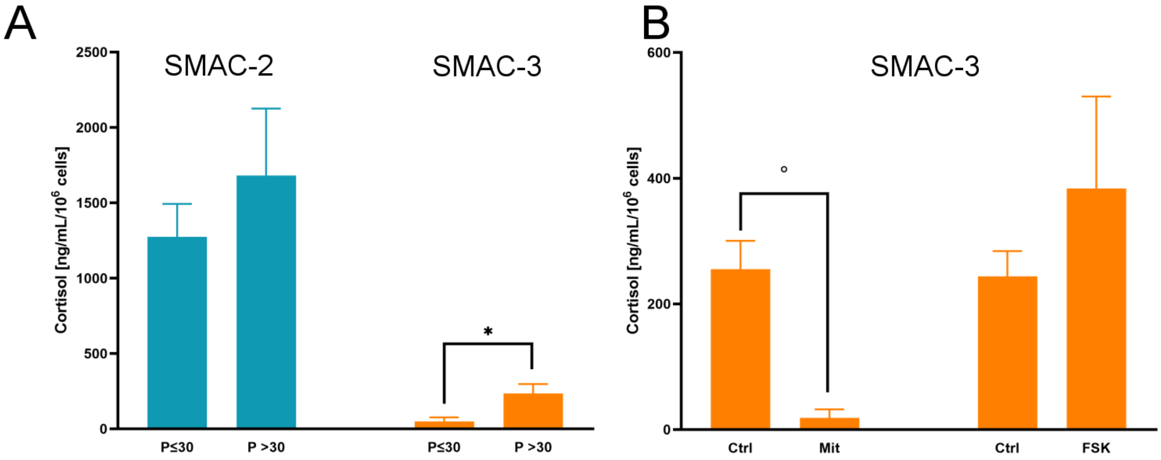


**Supplementary Figure 6. Effect on passages on cortisol secretion.** (A) Cortisol secretion of SMAC-2 and SMAC-3 at passages up to and after 30. Results are shown as ng/mL/$10^6$ cells of cortisol vs passages ≤ 30 ± SD. * $p < 0.0001$ vs passages ≤ 30. (B) Effect of mitotane and forskolin on cortisol secretion. Experiments were performed at passages > 30. Results are shown as ng/mL/106 cells of cortisol vs untreated cell ± SD.

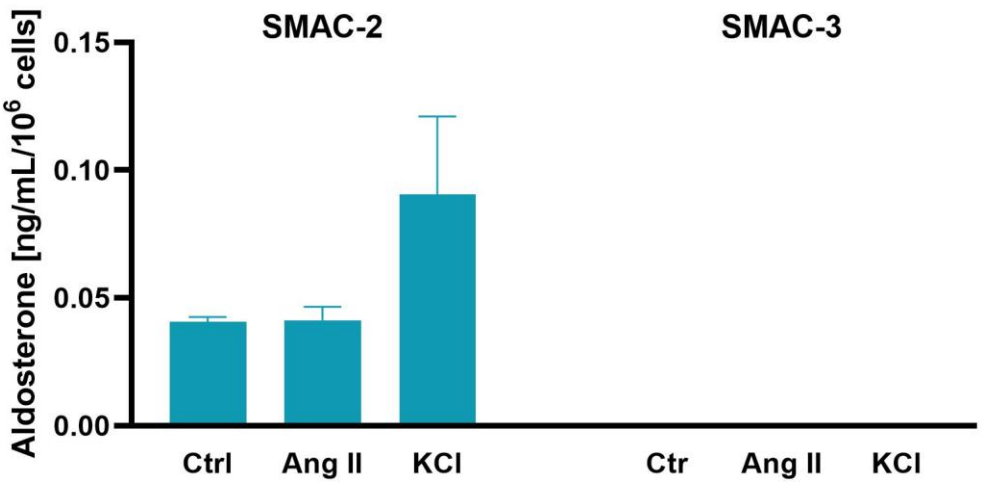


**Supplementary Figure 7. Effect of angiotensin II (Ang II) and potassium (KCl) on aldosterone secretion.** Results are shown as ng/mL/$10^6$ cells of aldosterone vs untreated cell ± SD.

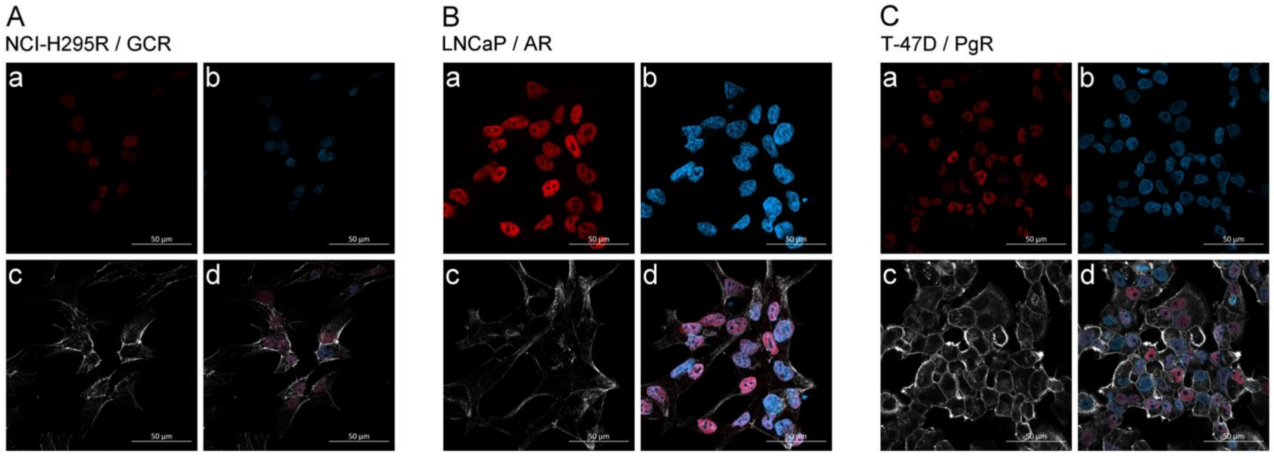


**Supplementary Figure 8. Intracellular steroid hormones receptors expression in positive controls cell lines.** Glucocorticoids receptor (GCR), androgens receptor (AR) and progesterone receptor (PgR) were evaluated in NCI-H295R, LNCaP and T-47D cells respectively. (aA) GCR, (aB) AR, (aC) PgR, (b) Hoechst 33258, (c) phalloidin and (d) merge. Images were acquired with a 40x objective. The scale bar of 50µm is automatically inserted by the software ZEN Blue.

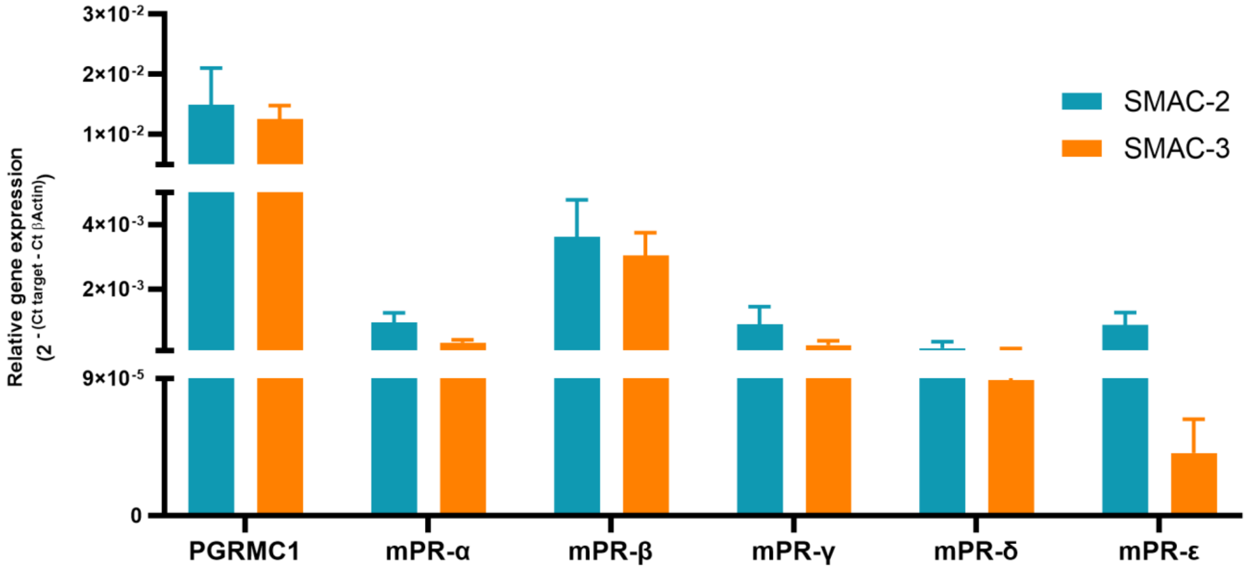


**Supplementary Figure 9. Gene expression of progesterone's membrane receptors.** Gene expressions were measured by q-RT-PCR using SYBR Green as fluorochrome. Results are shown as relative gene expression with β-actin as housekeeping gene ± SD.

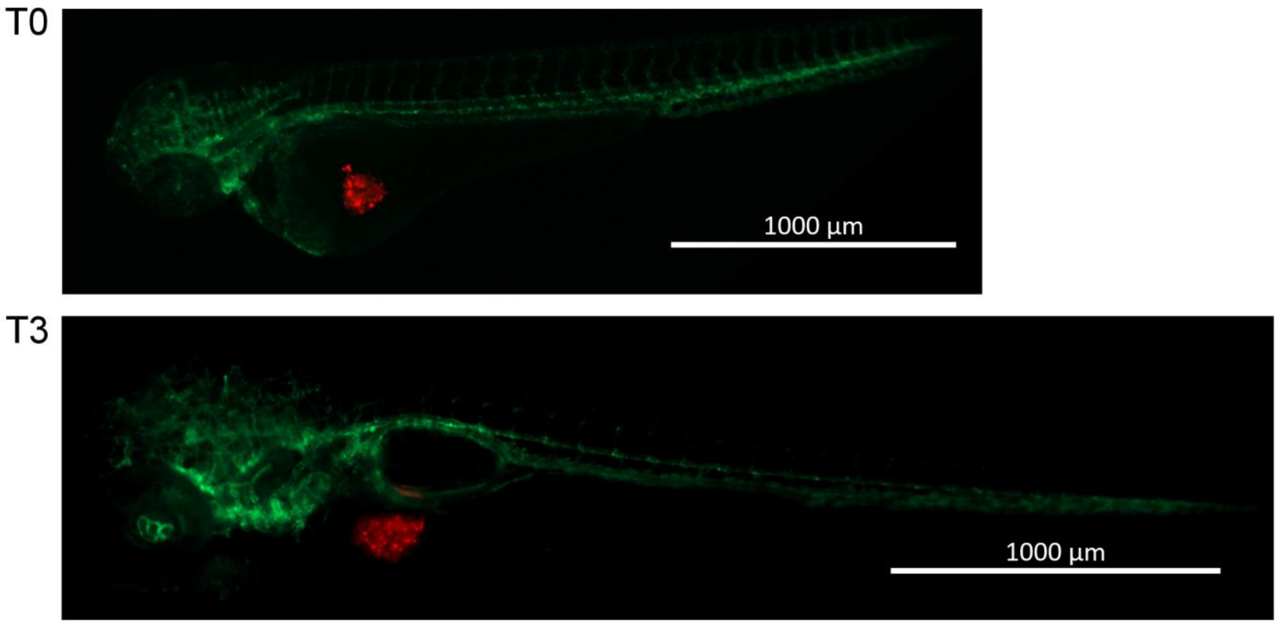


**Supplementary Figure 10. SMAC-3 cells xenograft in zebrafish embryos.** Representative images of 2 hours (T0) and 3 days (T3) post cells injection. SMAC-3 cells were labeled with a red fluorescent lipophilic dye while the embryo endothelium was labeled with a green fluorescent protein reporter driven by the kdrl promoter. Pictures of injected embryos were acquired using Zeiss Axiozoom V13 (Carl Zeiss AG) fluorescence microscope. The scale bar of 1000µm is automatically inserted by the software ZEN Blue

#### SUPPLEMENTARY TABLES

*Supplementary Table 1: Antibodies*

| Target | Characteristic | Company; cat. number | Dilution |
|---|---|---|---|
| **SF-1 (tissue)** | Rabbit | Abcam, Cambridge, United Kingdom; AB217317 | 1:500 |
| **Ki67** | Rabbit | Roche Diagnostics, Monza, Italy | prediluted |
| **SF-1 (cells)** | Rabbit | Cell Signaling Technologies, Milano, Italy; 12800 | 1:100 |
| **Ki67** | Rabbit | Cell Signaling Technologies | 1:400 |
| **β-catenin** | Rabbit | Cell Signaling Technologies; 8480 | 1:100 |
| **AR** | Rabbit | Cell Signaling Technologies | 1:200 |
| **GRC** | Rabbit | Cell Signaling Technologies | 1:100 |
| **PGR** | Rabbit | Cell Signaling Technologies | 1:500 |
| **Mouse** | Goat coniugate with Alexa Fluor 488 | Jackson ImmunoResearch, West Grove, USA; JI115545146 | 1:1000 |
| **Rabbit** | Goat coniugate with Alexa Fluor 594 | Jackson ImmunoResearch; JI111585144 | 1:1500 |
| **Phalloidin** | Coniugated with Alexa Fluor 647 | Invitrogen; A30107 | 1:400 |

*Supplementary Table 2: Prognostic and therapeutic targets*

| | | | | | | |
|---|---|---|---|---|---|---|
| AKT1 | CTNNB1 | GNA11 | KIT | NRAS | PPARG | SMAD4 |
| ALK | DDR2 | GNAQ | KRAS | NRG1 | PTEN | SMO |
| AR | EGFR | GNAS | MEK1 | NTRK1/2/3 | RAF1 | STK11 |
| BRAF | ERBB2/3/4 | HRAS | MET | PDGFRA | RB1 | TERT |
| CDK4 | ESR1 | IDH1/2 | MTOR | PIK3CA | RET | TP53 |
| CDKN2A | FGFR1/2/3/4 | KEAP1 | NF1 | POLE | ROS1 | TSC1 |

*Supplementary Table 3: Oligonucleotide sequences*

| Gene | Forward (5’ → 3’) | Reverse (5’ → 3’) |
|---|---|---|
| **SF-1** | CAGCCTGGATTTGAAGTTCC | TTCGATGAGCAGGTTGTTGC |
| **CYP11A1** | GAGATGGCACGCAACCTGAAC | CTTAGTGTCTCCTTGATGCTGGC |
| **CYP17A1** | GGCCCCATCTATTCGGTTCG | AGAGTCAGCGAAGGCGATAC |

| | | |
|---|---|---|
| **CYP21A2** | CGTGGTGCTGACCCGACTG | GGCTGCATCTTGAGGATGACAC |
| **CYP11B1** | TCCCGAGGGCCTCTAGGA | GGGACAAGGTCAGCAAGATCTT |
| **CYP11B2** | TCCAGGTGTGTTCAGTAGTTCC | GAAGCCATCTCTGAGGTCTGTG |
| **3βHSD2** | TGCCAGTCTTCATCTACACCAG | TTCCAGAGGCTCTTCTTCGTG |
| **CYP19A1** | AGGTGCTATTGGTCATCTGCTC | TGGTGGAATCGGGTCTTTATGG |
| **SULT2A1** | CCTCCAGCGGTGGCTACA | AATCGTCCGACATGATGATGAC |
| **GCR** | CCTACCCTGGTGTCACTGTT | CCTTTGCCCATTTCACTGCT |
| **AR** | CCTGGCTTCCGCAACTTACAC | GGACTTGTGCATGCGGTACTCA |
| **PgR** | CGCGCTCTACCCTGCACTC | TGAATCCGGCCT CAGGTAGTT |
| **PGRMC1** | CCATCAACGGCAAGGTGTTC | TCCAGCAAAGACCCCATACG |
| **MC2R** | CATGGGCTATCTCAAGCCAC | GAGATCTTCCTGGTGTGG |
| **β-Actin** | TCTTCCAGCCTTCCTTCCTG | CAATGCCAGGGTACATGGTG |

*Supplementary Table 4: reagent and condition for Sanger Sequencing*

| **Reagents and thermal profile** | | | | |
|---|---|---|---|---|
| **Reagents** | **for reaction** | **Temperature** | **Time** | **Cycles** |
| AmpliTaq Gold 360 Buffer, 10X | 2.50 µl | **95 °C** | 00:10:00 | |
| 25 mM Magnesium Chloride | 2.00 µl | **95 °C** | 00:00:30 | |
| dNTP mix | 1.00 µl | **61°C** *MSH2* (c.1222dup) | 00:00:30 | 30X |
| Primer Forward [0.4µM] | 1.00 µl | | | |
| Primer Reverse [0.4µM] | 1.00 µl | | | |
| 360 GC Enhancer | 0.50 µl | | | |
| AmpliTaq Gold 360 DNA Polymerase | 0.50 µl | | | |
| Nuclease free water | 11.00 µl | | | |
| DNA template (30ng) | 5.0 µl | | | |
| Extension | | **72°C** | 00:00:45 | |
| Final Extension | | **72 °C** | 00:07:00 | |
| Final Hold | | **4 °C** | Hold | |

*Supplementary Table: 5 Sanger sequencing PCR conditions and reagents*

| **Gene** | **Transcript** | **Exon** | **Coding** | **Forward 5’ ▯ 3’′** | **Reverse 5′ ▯3′** |
|---|---|---|---|---|---|
| *MSH2* | NM_000251.3 | 7 | c.1222dup | GTGGAAGCTTTTGTAGAAGATGCA | CTGAATGTGTCCTAAGAGTGAGTCA |
